# Protective Role of RIPK1 Scaffolding against HDV-Induced Hepatocyte cell death and the Significance of Cytokines in Mice

**DOI:** 10.1101/2023.10.15.562448

**Authors:** Gracián Camps, Sheila Maestro, Laura Torella, Carla Usai, Ana Aldaz, Cristina Olagüe, Africa Vales, Anne Montfort, Bruno Ségui, Lester Suarez, Rafael Aldabe, Gloria Gonzalez-Aseguinolaza

**Affiliations:** DNA & RNA Medicine Division, CIMA, University of Navarra, Instituto de Investigación Sanitaria de Navarra, IdisNA. Pamplona, Spain; Faculty of Pharmaceutical Sciences. Paul Sabatier University, Toulouse III INSERM UMR1037, Cancer Research Center of Toulouse (CRCT). Toulouse, France

**Keywords:** HDV, liver injury, RIPK1, Type I IFN, TNF-α, macrophages

## Abstract

Hepatitis delta virus (HDV) infection represents the most severe form of human viral hepatitis; however, the mechanisms underlying its pathology remain incompletely understood. We recently developed an HDV mouse model by injecting adeno-associated viral vectors (AAV) containing replication-competent HBV and HDV genomes. This model replicates many features of human infection, including liver injury. Notably, the extent of liver damage can be diminished with anti-TNF-α treatment. In the present study, we found that TNF-α is mainly produced by macrophages. Downstream of the TNF-α receptor (TNFR), the receptor-interacting serine/threonine-protein kinase 1 (RIPK1) serves as a cell fate regulator, playing roles in both cell survival and death pathways. In this study, we explored the function of RIPK1 and other host factors in HDV-induced cell death. We determined that the scaffolding function of RIPK1, and not its kinase activity, offers partial protection against HDV-induced apoptosis. A reduction in RIPK1 expression in hepatocytes through Cas9-mediated gene editing significantly intensifies HDV-induced damage. Contrary to our expectations, the protective effect of RIPK1 was not linked to TNF-α or macrophage activation, as their absence did not alter the extent of damage. Intriguingly, in the absence of RIPK1, macrophages confer a protective role. However, in animals unresponsive to type-I IFNs, RIPK1 downregulation did not exacerbate the damage, suggesting RIPK1’s role in shielding hepatocytes from type-I IFN-induced cell death. Interestingly, while the damage extent is similar between IFNAR KO and WT mice in terms of transaminase elevation, their cell death mechanisms differ. In conclusion, our findings reveal that HDV-induced type-I IFN production is central to inducing hepatocyte death, and RIPK1’s scaffolding function offers protective benefits. Thus, type-I IFN together with TNF-α, contribute to HDV-induced liver damage. These insights may guide the development of novel therapeutic strategies to mitigate HDV-induced liver damage and halt disease progression.

**Author summary:** Hepatitis D is the most aggressive form of viral hepatitis. Our manuscript underscores the complexity of HDV-induced liver damage, where both viral and host factors play significant roles. Previously, we demonstrated that pharmacological inhibition of TNF-α reduced HDV-induced liver damage. This result was corroborated in the present study using TNF-α-deficient mice. Moreover, we reported that the expression of the HDV antigen might have a cytotoxic effect, and HDV replication induces a strong activation of the innate immune system, accompanied by a substantial production of IFN-β. In this study, we discovered that RIPK1, a molecule described as a cell fate modulator acting downstream of TNF-α, plays a protective role during HDV replication. Contrary to our expectations, neither TNF-α nor macrophages, the primary producers of TNF-α, contributed to this protective effect. Instead, it seems type I IFN was involved. Interestingly, the role of type I IFN in HBV-induced liver damage has recently been proposed. Furthermore, our data reveals that several mechanisms of hepatocyte death are at play simultaneously during HDV replication, with apoptosis being one of them. Additional studies are needed to identify other mechanisms involved. Finally, these findings suggest that therapies targeting TNF-α and type-I IFN, or those increasing RIPK1 levels, might be effective in preventing or treating HDV-induced liver damage.

## Introduction

Hepatitis D virus (HDV) requires hepatitis B virus (HBV) for completion of its life cycle [1]. Coinfection with HBV and HDV is the most severe form of viral hepatitis, characterized by an increased risk of fulminant hepatitis, cirrhosis, liver decompensation, and hepatocellular carcinoma compared to HBV mono-infection [2]. It is estimated to affect 15-20 million individuals globally [3,4]. Despite the significant burden of this disease, the mechanisms underlying HDV-induced hepatocyte death and liver inflammation that conditions the severity of the disease and the rapid development of fibrosis are not fully understood [5]. HDV has previously been reported as not being directly cytotoxic to human hepatocytes. However, early studies showed that the expression of hepatitis D antigen (HDAg) resulted in significant cytotoxic changes in these cells [6]. We recently corroborated those results in mice overexpressing HDAgs in the liver of mice [7]. Although this can be seen as an artificial situation that doesn’t mirror the circumstances of natural viral infection, Ricco et al. found an association between HDV viral load and the severity of liver injury [8]. This brings up questions regarding the contribution of viral components to HDV-related pathology. Conversely, the role of the immune system in HDV pathogenesis is debated. Some studies emphasize the significance of HDV-specific adaptive immune responses [9,10], while others report weak or undetectable HDV-specific T cell responses. Moreover, there appears to be no correlation between the magnitude of HDV-specific CD4+ or CD8+ T cell responses and the clinical status of patients [11]. Interestingly, a recent study by Kefalakes et al. found a notable correlation between the size of both HDV-specific and non-specific CD8+ T cells expressing innate-like NKp30 (Natural Killer protein 30) and NKG2D (NK group 2D) receptors and the level of liver inflammation in HDV patients [12]. Furthermore, in vitro data showed that HDV infection of hepatic cell lines enhanced the cytotoxicity of CD8 T cells. This effect was independent of MHC class I-TCR interaction, but it did require activation of the innate immune response and type I interferon response [13]. Taken together, these results suggest that bystander activation of immune cells with cytotoxic capacity may contribute to liver pathology during HDV infection.

The lack of mouse models that accurately mimic human HDV infection has hindered progress in understanding the mechanisms underlying HDV-mediated liver damage. The available models, including immunodeficient mice with humanized liver or immunocompetent transgenic mice expressing the human sodium taurocholate cotransporting polypeptide (hNTCP) (receptor for HBV and HDV), do not fully replicate the characteristics of human HDV infection, particularly in terms of the induction of liver damage [14,15]. Recently, a HBV/HDV mouse model was developed using recombinant adenoassociated vectors (rAAV) to deliver HBV and HDV replication-competent genomes [7,16]. Interestingly, once HDV replication is established AAV genomes are hardly detectable, indicating that the rAAV acts as a HDV subtle and is not required for HDV-replication maintenance [16]. This model reproduces many of the features of HDV infection in humans, including liver inflammation and injury. Importantly, we found that liver injury can be partially ameliorated by TNF-α inhibition [7] which is in line with previous reports showing that TNF-α levels are higher in HDV-infected patients and that there is a close correlation between TNF-α and the severity of the disease [17]. TNF-α is an inflammatory cytokine recognized by TNF-α receptor 1 (TNFR1) on the cell surface, and downstream, the receptor-interacting serine/threonine-protein kinase 1 (RIPK1) acts as a cell fate regulator involved in both cell survival and cell death pathways [18]. The kinase activity of RIPK1 triggers cell death by apoptosis or necroptosis through the activation of caspase 8 or RIPK3-Mixed lineage kinase domain like protein (MLKL), respectively [19]. In line with this, necrostatin-1 (Nec1), an inhibitor of RIPK1 kinase activity, has been shown to have a protective effect in several models of acute liver injury, such as acetaminophen- and concanavalin-A-induced hepatocyte death [20,21]. However, RIPK1 may also play a protective role in hepatocytes [22–25]. In mice challenged with hepatotoxic agents such as lipopolysaccharides (LPS), poly(I:C) or murine hepatitis virus type 3 (MHV3), the absence of RIPK1 in hepatocytes resulted in significant elevation of liver transaminases and cell death [22–25]. Furthermore, it was demonstrated that the scaffolding function of RIPK1, but not its kinase activity, was responsible for this protective effect. Interestingly, the inhibition of TNF-α or depletion of macrophages resulted in amelioration of liver injury associated with the absence of RIPK1 [22,23].

In our current study, we determined the main cellular source of TNF-α in HDV-replicating mouse livers and the role of RIPK1 in HDV-induced liver injury. We found that the scaffolding function of RIPK1, but not its kinase activity, plays a protective role against HDV-induced liver damage. However, RIPK1 doesn’t block cell death induced by TNF-α and/or macrophages. Instead, it appears to target apoptosis induced by type-I IFN. Additionally, we also found that in the absence of RIPK1, macrophage depletion resulted in severe liver damage that can be fatal.

## Results

### Macrophages are the main source of TNF-α in the liver of AAV-HBV/HDV mice

In our previous study, we demonstrated that TNF-α plays a role in HDV-induced liver damage and its pharmacological inhibition showed a partial protective effect [7]. In order to determine which cells are responsible for TNF-α production we used RNA *in situ* hybridization (ISH) RNAscope® Fluorescent Multiplex Kit to analyze the localization of TNF-α mRNA and HDV genomic and antigenomic RNA in the liver of C57BL/6 wild type (wt) mice 21 days after AAV-HBV/HDV injection. First, we found a colocalization of HDV genomes and antigenomes in hepatocyte nuclei, indicating active replication of the virus (Fig 1A). We observed that most cells expressing TNF-α mRNA were HDV-negative (>80%) (Fig1A). Since HDV RNA expression and replication is restricted to hepatocytes by the use of a hepatocyte specific promoter, this result indicates that hepatocytes are not the main source of TNF-α. Previous studies have shown that liver macrophages are responsible for TNF-α production after various stimuli [19,20]. To analyze the role of this population, we combined TNF-α and HDV genome ISH with immunohistochemistry staining for the macrophage marker F4/80. As shown in Fig 1B-D, we found that most cells producing TNF-α are macrophages (Mean 65.9%), which in general are near HDV positive signals. Interestingly, 5 to 10% of macrophages expressing TNF-α contain HDV RNA, and all the macrophages positive for HDV are positive for TNF-α (Fig 1B). Then, we found 20-25% of cells expressing TNF-α that are F4/80 and HDV negative and a small number of HDV positive hepatocytes (Fig 1C).

**Fig. 1.**
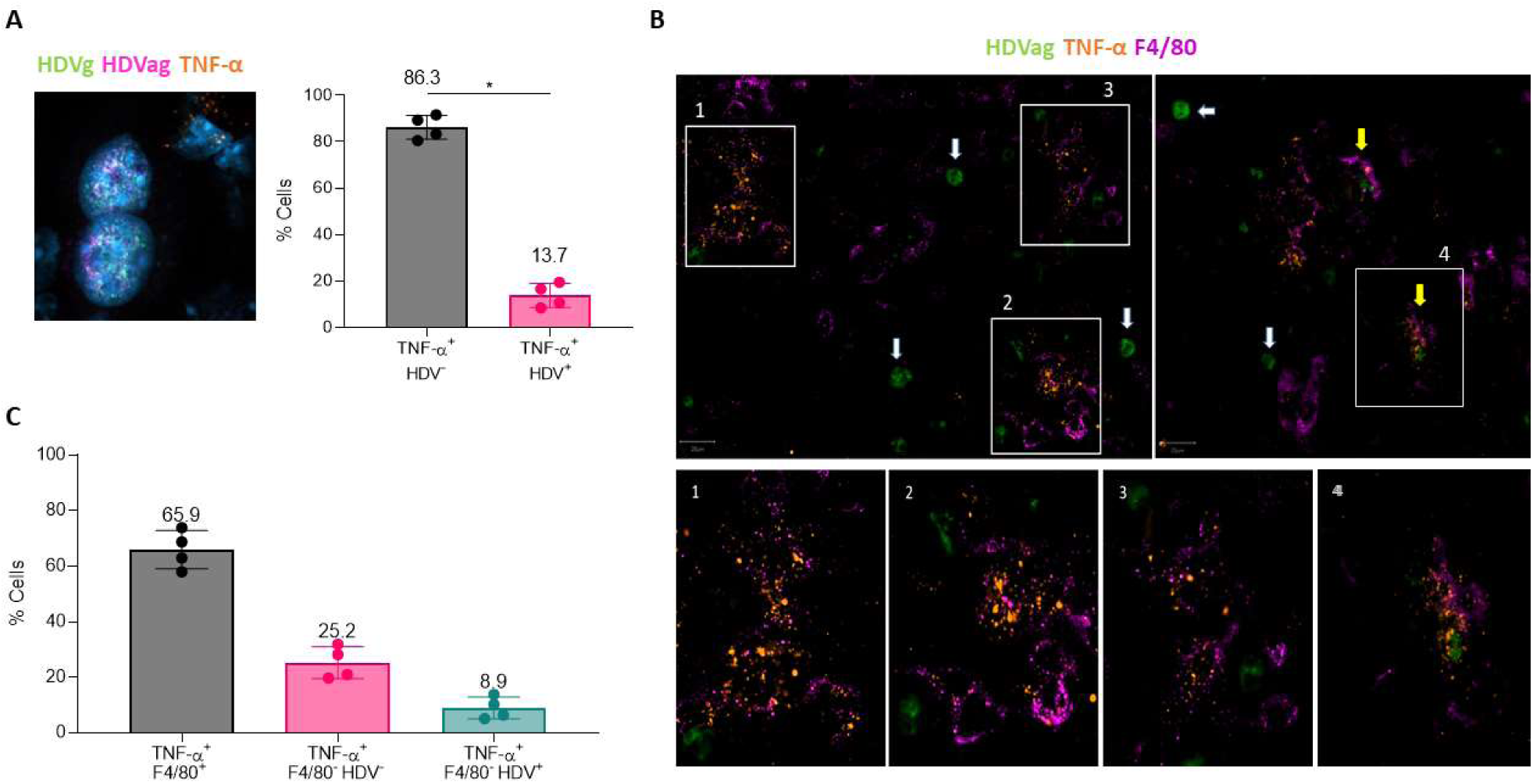
Macrophages are the main source of TNF-α production in AAV-HBV/HDV mouse livers. (A) TNF-α mRNA (orange) and HDV RNA genome (HDVg in green) and antigenome (HDVag in pink) distribution was analyzed by in situ hybridization (ISH) in the liver of C57BL/6 mice 21 days after receiving adenoassociated viral (AAV) vectors delivering both HBV and HDV genomes (HBV/HDV). An image showing the nuclei of two hepatocytes with HDV and HDVag signal and a close cell producing TNF-α. The percentage of TNF-α+ HDV+ and TNF-α+ HDV-was quantified using image J (mean ± SD, n=4). (B) Liver sections were analysed for TNF-α mRNA (green) and HDV RNA (orange) expression by in situ hybridization (ISH) followed by immunofluorescence (IF) to detect F4/80 protein (pink). Representative image of hybridized liver sections taking with the Vectra® PolarisTM Automated Imaging System are depicted in; white arrows pointed to HDV+ positive hepatocytes and yellow arrows pointed to macrophages expressing TNF-α, some of them are shown at a larger magnification in 1-4 images including macrophages containing HDV RNA (4). (C) The percentage of TNF-α+ F4/80+, TNF-α+ F4/80+ HDV- and TNF-α+ F4/80+ HDV+ cells was quantified using image J. Individual data points and mean values ± standard deviations are plotted. Statistical differences were determined by Mann-Whitney test p <0.05 (*).

### RIPK1 scaffolding function, but not its kinase activity, is critical for the survival of HDV-infected hepatocytes

Several studies have shown the key role of RIPK1 in maintaining liver homeostasis under conditions of macrophage activation and TNF-α production [22–25]. To assess the role of RIPK1 in HDV-mediated liver injury, we developed a CRISPR/Cas9 gene editing system that specifically knockdown RIPK1 expression in hepatocytes. We achieve this by expressing the Cas9 protein under the control of a liver specific promoter (S1A Fig) that was delivered using a rAAV vector with hepatic tropism. Two different guide (g) RNAs were designed (RIPK1g1 and RIPK1g2) and their editing efficacy was tested *in vivo* by analyzing RIPK1 protein expression by western blot. As shown in S1B Fig, RIPK1 protein expression was significantly reduced with RIPK1g2 and selected for further studies (the AAV vector containing Cas9 and RIPK1g2 was named RIPK1edit). To determine the functional effects of RIPK1 downregulation, wt mice were treated with RIPK1edit and their susceptibility to LPS-induced liver injury and apoptotic cell death, which have been shown to be exacerbated in the absence of RIPK1 [22], was evaluated as described in S1C Fig. We found that RIPK1edit-treated animals showed a significantly higher susceptibility to LPS-induced liver damage compared to control animals, as shown by higher levels of transaminases and frequency of apoptotic cells identified by the expression of activated Caspase-3 (a-Casp3) (S1D-F Fig), demonstrating the functional impact of RIPK1edit in the liver.

To investigate the role of RIPK1 in HDV-induced liver injury, wt mice were administered with RIPK1edit or a rAAV expressing Cas9 but without guide (control). Thirty hours later (before the development of anti-AAV neutralizing antibodies), each group was divided into two subgroups that received AAV-HBV/HDV (HBV/HDV) or AAV-HBV (HBV), respectively (Fig 2A). Transaminase levels were evaluated weekly up to day 21, when the animals were sacrificed (Fig 2A). As expected from our previous data [7,16] no liver damage was observed in control mice receiving HBV while a significant increase in transaminase levels was observed at day 14 and 21 in control HBV/HDV animals. RIPK1edit has no effect over HBV induced liver injury; however, it has an important impact over HBV/HDV-induced liver injury since transaminase levels were significantly higher in RIPK1edit-HBV/HDV mice in comparison to control-HBV/HDV mice 14 and 21 days-post-infection (dpi) (Fig 2B). At sacrifice, the expression of a-Casp3 and HDV antigen (HDAg) was analyzed by IHC and HDV genome levels were measured by RT-qPCR. We observed a higher percentage of a-Casp3 positive (apoptotic) hepatocytes in RIPK1-edit mice receiving HBV/HDV compared to control HBV/HDV mice; while no apoptotic cells were detected in the two groups of mice receiving HBV alone (Fig 2C). Moreover, there was a direct correlation between liver damage and the number of apoptotic hepatocytes in the two groups of mice receiving HBV/HDV (S2A Fig). Interestingly, while there were no major differences in HDV viremia between control and RIPK1edit mice (Fig 2D), we observed that the number of cells expressing HDAg (Fig 2E) and the HDV genome levels (data not shown) at day 21 were significantly lower in RIPK1edit mice, likely due to increased hepatocyte death.

**Fig. 2.**
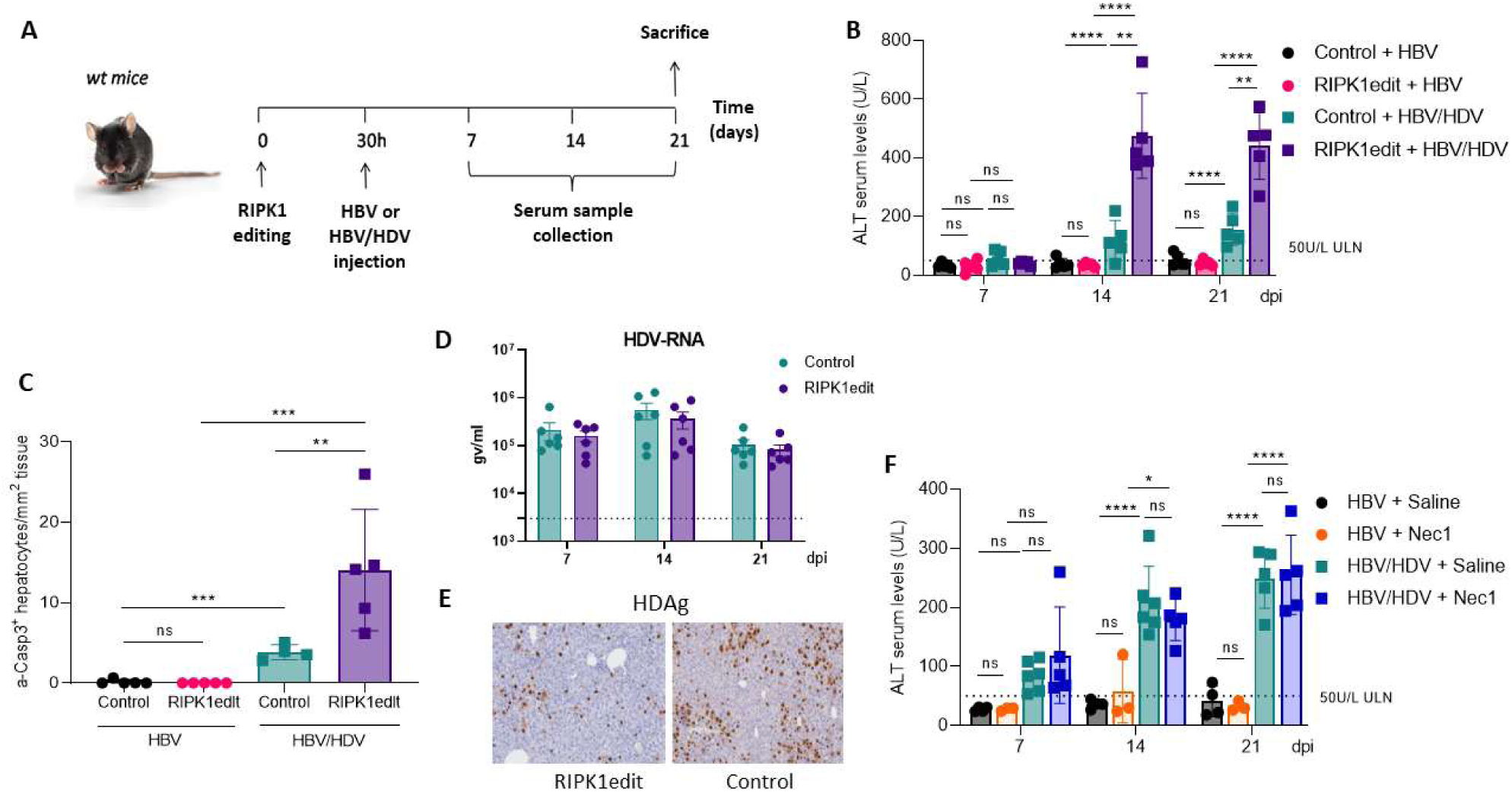
RIPK1 scaffolding function partially prevents hepatocellular apoptosis in HBV/HDV mice. (A) Schematic representation of the experimental procedure, 6-8-week-old C57BL/6 wt mice were intravenously injected with 10^11^ genome copies (gc) of AAV-SaCas9-RIPK1g2 (RIPK1edit) or an AAV expressing SaCas9 without guide (control) and after 30 hours they were administered with 5 x 10^10^ gc AAV-HBV (HBV) or AAV-HBV/HDV (HBV/HDV). Peripheral blood was extracted at 7, 14 and 21 days post-injection (dpi). Liver damage was analysed by B) quantification of serum ALT levels (U/L) and (C) quantification of a-Casp3+ hepatocytes/area after Immunohistochemistry (IHC). (D) HDV viremia in circulation was determined in control and RIPK1edit mice 7, 14 and 21 dpi and (E) IHC analysis against HDAgs was performed at 21 dpi in the liver sections of both groups. (F) Analysis of serum ALT levels in wt mice daily treated with Nec1 or saline was performed 7, 14 and 21 days after administration of HBV or HBV/HDV. Individual data points and mean values ± standard deviations are shown. Statistical analysis was performed by one-way ANOVA followed by Bonferroni multiple-comparison test. p <0.05 (*), p <0.01 (**), p <0.001 (***), p <0.0001 (****) and ns = non-significant.

The experiment was repeated in C57BL/6 Rag1 deficient mice to evaluate the potential role of the adaptive immune system in the exacerbation of liver damage upon RIPK1 downregulation. As shown in S2B-D Fig, in the absence of B, T, and NKT cells, HBV/HDV-induced liver damage was also exacerbated after RIPK1 knockdown as shown by significant higher transaminase levels (S2B Fig) as well as a higher number of apoptotic hepatocytes (S2C Fig) and both parameters showed a direct correlation (S2D Fig).

To determine if the exacerbation of HBV/HDV-induced liver damage observed in RIPK1edit mice was related to RIPK1 kinase activity, HBV/HDV injected animals were treated daily with a dose of 2.5 mg/kg Nec1, and HBV injected mice were used as controls. All animals were bled weekly and sacrificed at 21 dpi. The efficacy of Nec-1 treatment was evidenced by a significant reduction of MLKL expression levels in the liver of mice (S2E Fig). As shown in Figure 2E, transaminase levels in HBV/HDV injected mice were not affected by Nec-1 treatment. Thus, this result indicates that RIPK1 kinase activity appears to play no role in the protective effect of RIPK1 during HDV replication.

In summary, our data suggest that the scaffolding function of RIPK1 is critical for the survival of HDV-infected hepatocytes and that the increased liver damage in RIPK1-edited animals is independent of the adaptive immune response.

### Macrophages and TNF-α are not involved in the exacerbation of HDV-mediated liver damage in the absence of RIPK1

Previous studies have shown that, in the absence of RIPK1, liver damage induced by the administration of different agents to mice is attenuated by macrophage depletion or anti-TNF-α treatment [22,23]. Here, we examined the role of both in the increase of liver damage observed in HBV/HDV-infected mice after RIPK1 downregulation. As a first step, we analyzed the presence of macrophages in the livers of mice 21 days after HBV/HDV or HBV injection in RIPK1edit and control mice by F4/80 IHC. We found a significant increase in liver macrophages in HBV/HDV mice in comparison to HBV mice, that were further increased when mice were pretreated with RIPK1edit (Fig 3A, B). In order to determine the role of the macrophage population in the exacerbation of liver damage due to the absence of RIPK1, they were removed using chlodronate-loaded liposomes (CLL) and liver injury was evaluated. CLL was administered i.v. 2 days before HBV or HBV/HDV injection and then every 4 days up to day 21 pi. Macrophage-depletion was confirmed by F4/80 IHC (S3A-B Fig). In control animals (AAV expressing Cas9 but no guide) macrophage depletion resulted in a slight reduction of liver damage after HBV/HDV injection at 21 dpi (Fig 3C) and in a significant reduction of TNF-α expression (S3C Fig). However, in RIPK1edit the administration of CLL had a severe detrimental effect with a highly significant increase in transaminase levels in comparison to control mice (Fig 3C). Interestingly, the number of apoptotic hepatocytes in RIPK1-edited and CLL-treated mice was lower than in RIPK1-edited non CLL-treated mice (Fig 3D), and liver histology revealed the presence of necrotic areas (Fig 3E), indicating activation of a death mechanism other than apoptosis.

**Fig. 3.**
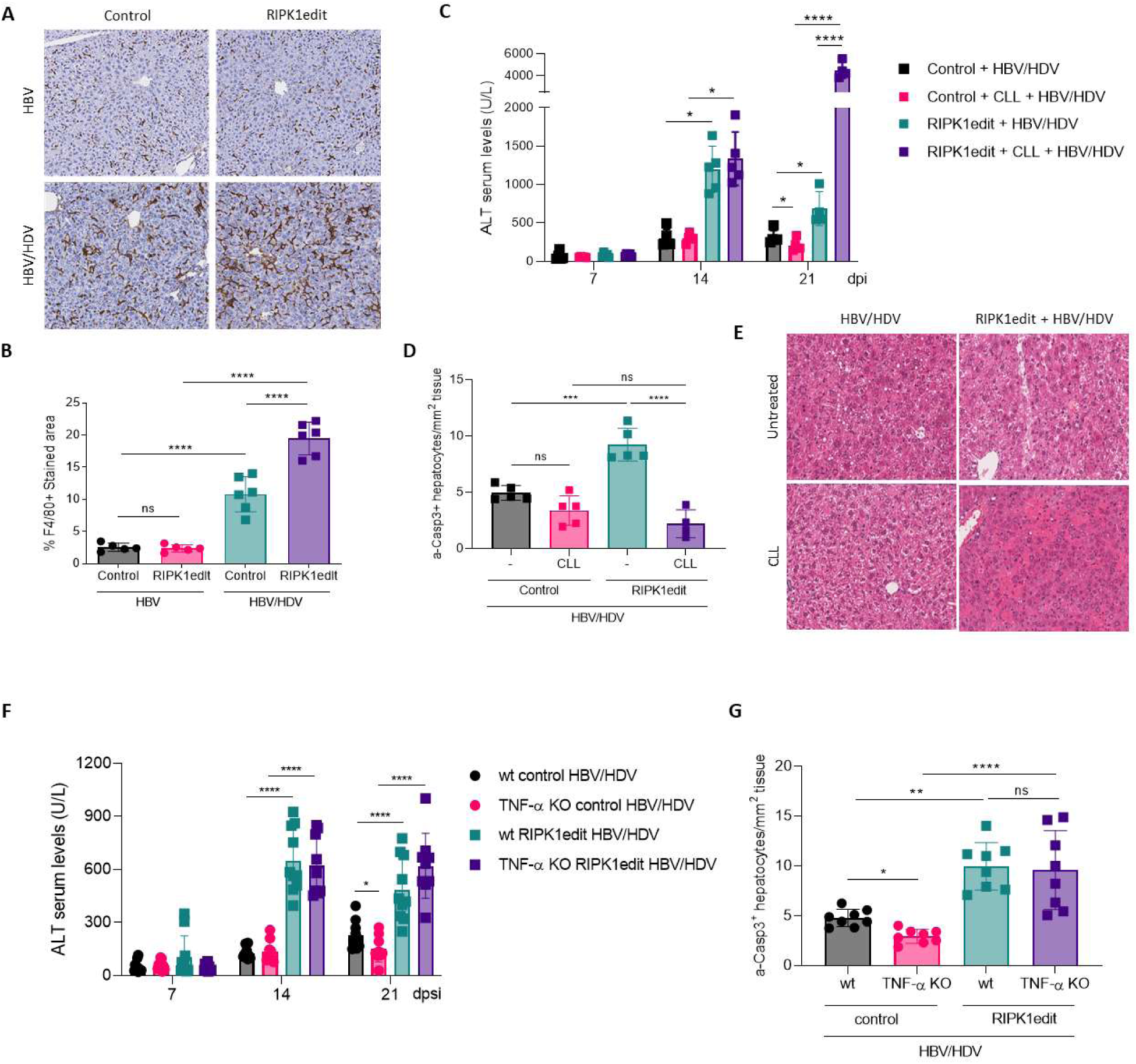
The exacerbation of liver damage in RIPK1-edited mice upon HBV/HDV co-injection is not mediated by macrophages or TNF-α. The presence of macrophages in the livers of control and RIPK1edit wt mice was analyzed 21 dpi of HBV or AAV-HBV/HDV by F4/80 immunohistochemistry (IHC). (A) Representative IHC images of F4/80-stained livers are shown (200x). (B) Quantification of F4/80 IHC stained area (mean ± SD). Significant differences were determined by one-way ANOVA followed by Bonferroni multiple-comparison test. (C-E) 6-8-week-old C57BL/6 wt mice were treated as described in Fig.2A and for macrophage depletion, animals were intravenously injected with clodronate-loaded liposomes (CLL) every 4 days starting the day before HBV/HDV injection up to day 20. Liver damage was analyzed by (C) determination of serum ALT levels (U/L) (D) quantification of a-Casp3+ hepatocytes/area (Individual data points and mean values ± SD are shown) and (D) liver histology analysis on H&E-stained sections, representative images are shown. Statistical differences were determined by two-way (D) or one-way (E) ANOVA followed by Bonferroni multiple-comparison test. (F-G) 6-8-week-old C57BL/6 wt and TNF-α KO mice were treated as described in Fig.2A. Liver damage was analyzed by (F) determination of serum ALT levels (U/L) and (G) quantification a-Casp3+ hepatocytes. Individual data points and mean values ± standard deviations are shown. Statistical differences were determined by two-way (F) or one-way (G) ANOVA followed by Bonferroni multiple-comparison test. P <0.05 (*), p <0.01 (**), p<0.001 (***), p <0.0001 (****) and ns = non-significant.

Next, we assessed the role of TNF-α in the increased liver damage in RIPK1-edited mice due to HBVHDV by employing C57BL/6 mice lacking TNF-α. TNF-α KO mice receiving HBV/HDV showed lower transaminase levels than wt animals 21 dpi (Fig 3F), in line with our previous published data [7]. However, transaminase levels and the number of apoptotic hepatocytes in RIPK1-edited mice were similar in TNF-α KO and wt mice (Fig. 3F, G), indicating that the exacerbation of liver damage observed in RIPK1edit mice is not related to TNF-α production.

In conclusion, neither macrophages nor TNF-α play a major role in the increased liver damage observed in HBV/HDV-infected mice in the absence of RIPK1. Instead, our data suggest that macrophages play a protective role during HBV/HDV infection in RIPK1-knockdown mice, in which apoptotic cell signaling is intensified.

### HDV Ag-mediated hepatocyte apoptosis it is not affected by the absence of RIPK1

In our previous study we have shown that HDV small and large antigens (S-HDAg and L-HDAg), in particular the S-HDAg, are also involved in the induction of hepatocyte death [7]. In order to determine if RIPK1 plays a role in protecting hepatocytes form HDV-Ag-induced cell death, RIPK1 edited animals received a dose of an AAV expressing S-HDAg and transaminase levels were analyzed 7 and 14 days after vector administration (the time points we have previously determined transaminase levels were elevated [7]. As shown in Fig 4A, no differences in transaminase levels were observed between control and RIPK1 edited animals. Furthermore, the levels of apoptotic cells although higher than in control animals were very low and not altered by RIPK1 editing (Fig 4B), suggesting that the main mechanism of HDAg-induced cell death is not apoptosis and RIPK1 played no role. Importantly, no major differences in the S-HDAg expression levels were observed between the two groups (Fig 4C).

**Fig. 4.**
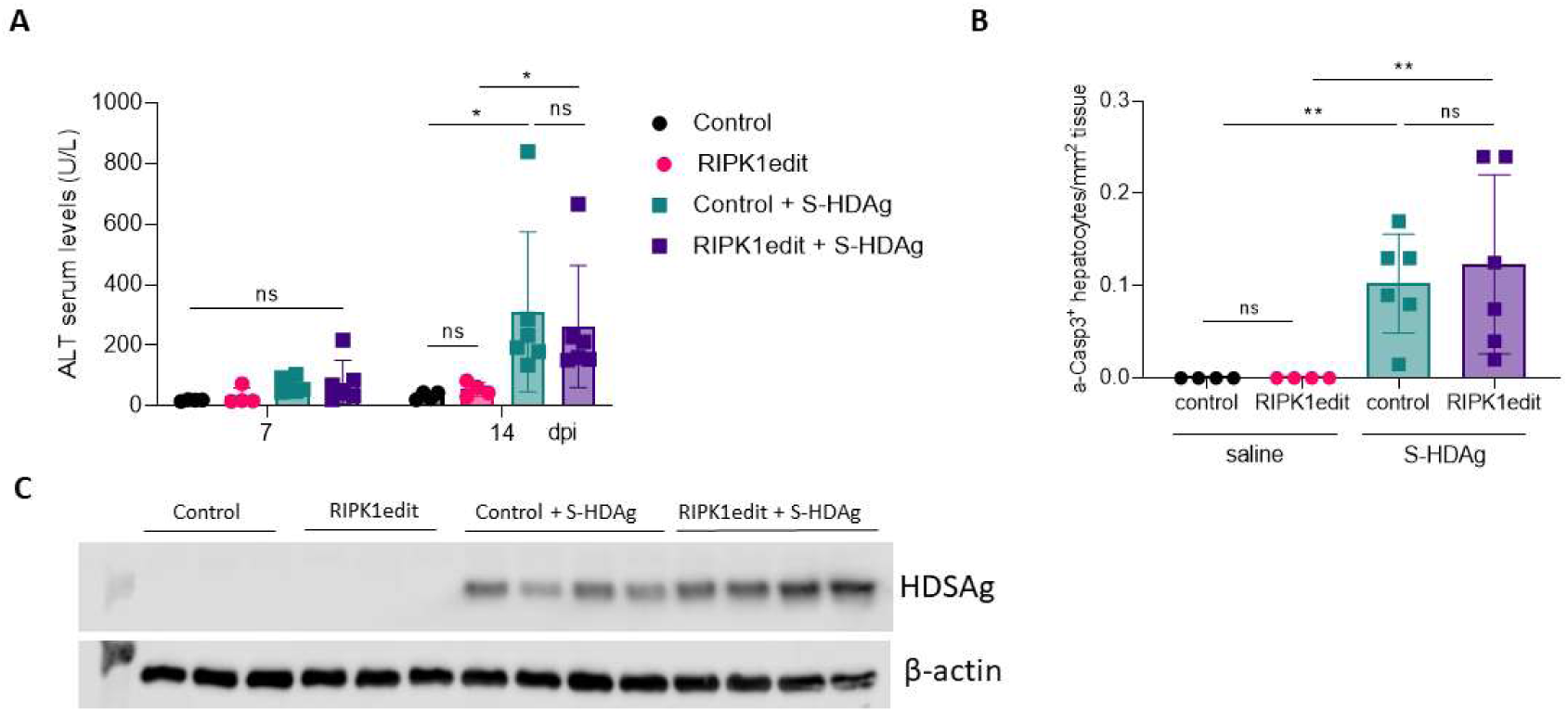
RIPK1 downregulation has no effect over HDAg-induced liver damage. 6-8-week-old C57BL/6 wt were injected with AAV-Cas9 control or RIPK1 edit and 30 hours later both groups were injected with an AAV expressing HDV-Small Antigen (S-HDAg). Liver damage was analyzed by (A) determination of serum ALT levels (U/L) at 7 and 14 days post injection (dpi) and (B) quantification of a-Casp3+ hepatocytes/area at 14 dpi. (C) Western blot analysis to determine S-HDAg expression was performed at 14 dpi in the liver extracts. Individual data points and mean values ± standard deviations are shown in A and B. Statistical differences were determined by two-way ANOVA.

### RIPK1 downregulation has no effect in mice lacking type I IFN receptor

We have previously shown that HDV replication, using the AAV-HBV/HDV animal model, leads to the production of high levels of type-I IFN via the MAVS signaling cascade [16]. This has also been confirmed *in vitro* using human cells [13,26]. We also reported no differences in the magnitude of HDV-induce liver damage between C57BL/6 mice lacking the IFNα/β receptor (IFNα/βR KO) and wt mice, thus we concluded that type I IFN played no role on HDV-induced liver damage [7]. However, we did not investigate the cell death mechanism in those animals.

Here to determine if in the absence of RIPK1, type-I IFN has a role in the exacerbation of liver damage induce by HBV/HDV replication, we treated IFNα/βR KO and wt control with RIPK1edit followed by HBV/HDV. As shown in Figure 5, and as previously reported [7], there were no significant differences in transaminase levels between wt and IFNα/βR KO mice after HBV/HDV injection. However, while RIPK1 knockdown had a significant impact on the liver damage in wt animals, there were no differences in transaminase levels in IFNα/βR mice regardless of RIPK1 expression (Fig 5A). Interestingly, the number of a-casp3 was lower in IFNα/βR than in wt, indicating differences in the mechanism of HDV-induced cell death in the absence of response to type-I IFN (Fig 5B). In addition, the analysis of HDAg expression revealed that the number of positive cells were very similar in control wt and IFNα/βR KO mice. However, whereas RIPK1 downregulation resulted in a decrease of HDAg expressing cells in wt mice, this decrease was not observed in IFNα/βR KO animals, which correlates with the absence of disease exacerbation in these animals (Fig 5C,D). Collectively, these results suggest that when response to type-I IFN is intact HDV-induces apoptotic cell death and RIPK1 plays a protective role but in its absence other cell death mechanisms seem to come into play.

**Fig. 5.**
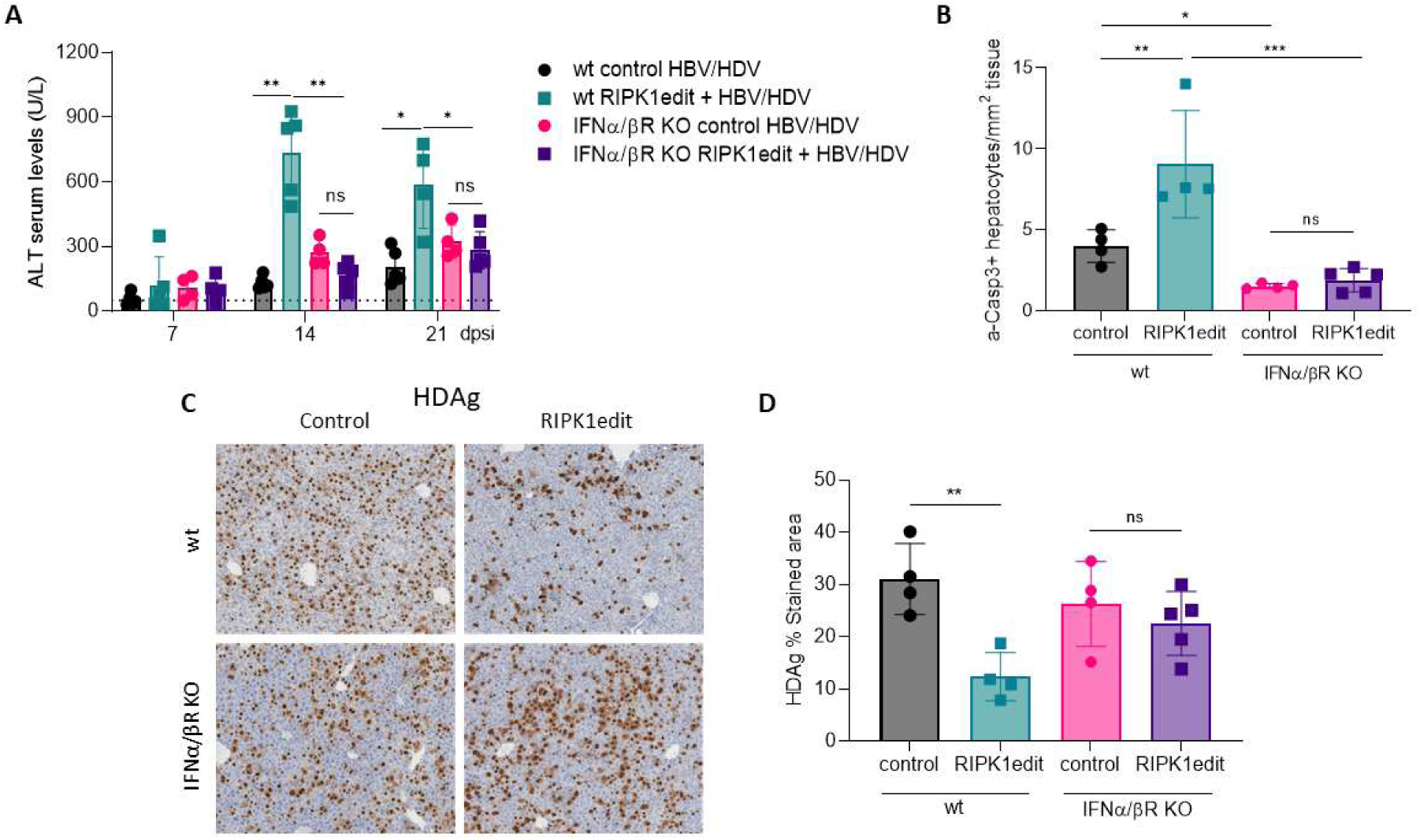
RIPK1 downregulation has no effect over HDV-induced liver damage in the absence of type-I IFN response. 6-8-week-old C57BL/6 wt and IFN-α/βR KO mice were treated as described in Fig.2A. Liver damage was analyzed by (A) determination of serum ALT levels (U/L) at 7, 14, and 21 days post injection (dpi) and (B) quantification of a-Casp3+ hepatocytes/area at 21 dpi. (C) IHC against HDAgs was performed at 21 dpi in the liver sections of control and RIPK1edit mice injected with the HBV/HDV vectors. (D) Quantification of HDAg expression in the aforementioned IHC images for each mouse. Individual data points and mean values ± standard deviations are shown. Statistical differences were determined by two-way ANOVA (A) or one-way ANOVA (B and D) followed by Bonferroni multiple-comparison test. p <0.01 (**), p <0.001 (***), p <0.0001 (****) and ns = non-significant.

## DISCUSSION

Cell death plays a central role in the resolution and progression of disease. During viral infection, controlled cell death can eliminate pathogens, but it can also cause uncontrolled organ damage [27]. HDV infection is the most aggressive form of viral hepatitis infection, however the mechanism involved in HDV-induced cellular death has not yet been fully understood. While apoptosis is a prevalent form of cell death in hepatocytes, other mechanisms such as necroptosis, ferroptosis, and autophagy-mediated cell death can also occur [28]. Liver parenchymal cells can activate different cell death pathways simultaneously, depending on the circumstances. The type of cell death that ultimately occurs is determined by factors such as the intensity and nature of the signal, the cellular context, and the presence of inhibitors or activators of specific cell death pathways [28]. Thus, understanding the complexity of the underlying mechanisms and the interplay between different pathways is crucial for developing effective strategies for treating liver diseases.

Previously, we showed that replication of HDV and HBV in mouse hepatocytes after AAV-mediated genome delivery induced significant liver damage, accompanied by the production of various cytokines including TNF-α and IFN-β [7]. In addition, we found that pharmacological inhibition of TNF-α resulted in a significant reduction in HDV-mediated liver damage [7]. These results have been corroborated in the present study using TNF-α deficient mice in which we observed a significant reduction, but not elimination, of HDV-induced liver injury. Here, we determined that TNF-α is primarily produced by activated macrophages. Consistent with this observation, depletion of macrophages significantly reduced TNF-α production and also partially attenuated HDV-induced liver damage. Our results suggest that HDV-induced hepatocyte-derived signals likely activate macrophages, which are responsible for producing this cytokine. Interestingly, we also found a non-negligible percentage of macrophages that were positive for HDV RNA. Since initial HDV RNA synthesis in our system is controlled by a liver-specific promoter, and mouse cells cannot be directly infected by circulating HDV infectious particles, this finding suggests that HDV RNA is transferred from hepatocytes to macrophages through an alternative mechanism and not by AAV-HDV-mediated transduction or HDV infection. It has recently been described that HDV can efficiently spread without envelopment through the proliferation of infected cells [29], so one possibility is that HDV-positive hepatocytes proliferate and spread HDV genomes to other cells, including macrophages. Another potential explanation is that macrophages uptake extracellular vesicles (EVs) produced by hepatocytes harboring HDV replication. The formation of EVs from HDV- and HBV-infected hepatocytes has been reported by other groups [30,31]. EVs can be taken up by monocytes, macrophages, and dendritic cells, inducing their activation [32], in fact, all the macrophages in which HDV was detected also expressed TNF-α.

Different studies have shown that TNF-α produced by liver macrophages plays a key role in both acute and chronic liver disease [22,23]. As described, TNF-α is recognized by TNFR1, which interacts with RIPK1, a protein with kinase activity that has emerged as a central molecular switch in controlling the balance between cell survival and cell death [18]. Depending on the type of insult, RIPK1 can promote cell survival, apoptosis, or necroptosis [19–25]. To study the potential role of RIPK1 in HDV-induced cell death, we developed a hepatocyte-specific gene-editing system using CRISPR-Cas9 delivered by an hepatotropic rAAV vector. Using this in vivo editing system, we found that downregulation of RIPK1 expression resulted in a significant increase in liver damage after HBV/HDV injection that was not observed when HBV was administered alone. This suggests that RIPK1 plays a protective role during HDV infection. However, inhibiting RIPK1 kinase activity with Nec-1 showed that this protection is not related to its kinase activity. This suggests it is most likely related to its scaffolding function, as previously demonstrated in other contexts [25]. Furthermore, the exacerbation of liver damage in the absence of RIPK1 was accompanied by an increase in the number of apoptotic cells. Although T cells have been suggested to be involved in HDV-induced liver damage [9,10], we found that immunodeficient mice lacking T cells had a similar magnitude of liver injury to wt mice and the downregulation of RIPK1 has a similar effect, indicating that other mechanisms independent of an uncontrolled T cell immune response against HDV-infected cells are involved in the pathology.

Interestingly, different studies have shown that the death of RIPK1-deficient hepatocytes induced by various agents (such as MHV3 infection, Poly(I:C), concanavalin A, LPS, or α-galactosylceramide) is mediated by TNF-α primarily produced by liver macrophages [20–25]. However, to our surprise, the absence of TNF-α or the elimination of macrophages did not alleviate HDV-induced liver damage in the absence of RIPK1. In fact, the depletion of liver macrophages had a clear detrimental impact. RIPK1 gene-edited mice in which macrophages were depleted showed a dramatic increase in hepatic injury after HBV/HDV administration, and the animals were nearly moribund at the end of the study. Histologically, we observed the formation of large necroinflammatory foci, which probably reflects that apoptotic hepatocytes were not removed by macrophages, a process that is crucial for the maintenance of liver health and homeostasis [34]. These results support the evaluation of macrophages as a potential cell-based therapy for the control of HDV-induced liver damage, a strategy that is currently being evaluated in clinical trials for the treatment of liver cirrhosis [35].

Previously, we and others have shown that HDV antigens may play a role in HDV-induced liver damage [6,7]. Thus, we explored whether RIPK1 was involved in reducing the toxic effects produced by HDAg, but we observed no differences in the magnitude of liver damage induced by HDAg overexpression in the presence or absence of RIPK1.

Finally, we investigated the role of type-I IFN response in the exacerbation of liver damage in the absence of RIPK1. We have previously demonstrated in the HBV/HDV mouse model that HDV replication induced MAVS-mediated IFN-β production [16]. This result was corroborated in human cells showing that MDA5 was involved in the detection of HDV RNA which results in the production of type I IFN [13,26]. Interestingly, RIPK1 downregulation in IFNα/βR KO mice had no effect on the liver damage induced by HDV. Specifically, the magnitude of this damage, gauged by transaminase levels, remained consistent regardless of the presence or absence of RIPK1. Furthermore, in line with our previous results, we observed no differences in the severity of HDV-induced liver injury between wt and IFNα/βR KO mice. However, when examining the number of apoptotic cells, we found a significantly lower count in IFNα/βR KO mice compared to wt mice, regardless of the presence or absence of RIPK1. This suggests that in the absence of a response to type I IFN, the mechanism of HDV-induced cell death largely differs, though not entirely, from that in wt mice.

These results might suggest that in wt mice, type I IFN may be involved in HDV-induced apoptosis, and RIPK1 plays a protective role. Recently, it has been shown that type I IFN signaling induces cell death in HBV-hepatocytes by suppressing the unfolded protein response [36]. Furthermore, constitutive activation of MDA5 has been shown to lead to cell death [37], and our previous data showed sustained activation MAVS-MDA5 activation in AAV-HDV mice thus it is not unlikely that type-I IFN is involved in HDV-induced liver damage [16]. Nonetheless, there is limited information regarding the interplay between type I IFN and RIPK1 and additional studies are needed.

In summary, our data indicate that HDV-induced cell death is complex, with both viral and host factors involved. Explaining why eliminating just one of these factors has only a partial effect on the extent of the damage. We found that type I IFN, along with TNF-α, are responsible for HDV-induced liver damage. It appears that various cell death modes coexist in the HDV-affected liver, a phenomenon recently termed “PANoptosis.” PANoptosis describes a novel cell death routine where various cell death modes are engaged simultaneously, in some cases in response to pathogen’s infection [38]. Remarkably, it has been shown that RIPK1 regulates Yersinia-Induced PANoptosis [39]. Further research is required to understand the interplay between different cell death pathways and the potential role of PANoptosis in HDV-induced liver disease. This knowledge will enhance our understanding of the intricacies and connections in HDV-induced pathogenesis and assist in the development of therapeutics.

## Material and methods

### Animals and treatments

C57BL/6 mice were purchased from Harlan Laboratories (Barcelona, Spain). Rag1-(Rag1 KO), IFNα/β receptor-(IFNα/βR KO) and TNF-α (TNF-α KO) deficient mice, all of them on a C57BL/6 background, were bred and maintained at the animal facility of Cima-Universidad de Navarra. Six- to eight-week-old male mice were used in all experiments. Mice were kept under controlled temperature, light, and pathogen-free conditions. Blood collection was performed by submandibular bleeding, and serum samples were obtained after centrifugation of total blood. Animals were euthanized by cervical dislocation after being anesthetized at the indicated time points. Liver samples were collected for histological analysis and for protein extraction. The HBV and HBV/HDV mouse model were generated by the intravenous (iv) administration of AAV-HBV or AAV-HBV+AAV-HDV, respectively as previously described [16] RIPK1 gene edition was performed by iv injection of AAV carrying liver specific CRISPR-Cas9 editing systems [40]. Lipopolysaccharide (LPS) (Sigma-Aldrich) were administered intraperitoneally at 0.4 μg/g and 5 μg/g of body weight, respectively, in a volume of 100 μl. Necrostatin-1 (Nec-1) was administered intraperitoneally at a dose of 2,5 mg/kg mg/kg. Macrophage depletion was achieved by intravenous (i.v.) administration of 100 μl clodronate-loaded liposomes (Clodlip BV). The experimental design was approved by the Ethics Committee for Animal Testing of the University of Navarra.

### Multiplex Fluorescence RNA In situ Hybridization (ISH) and ISH in combination with F4/80 immunofluorescence (IF)

In situ Hybridization (ISH) was performed with the ACD RNAscope® Fluorescent Multiplex Kit (Advanced Cell Diagnostics, USA) in liver sections. Firstly, livers were cryopreserved in an OCT cryomold and were cut in 10 μm-sections. Then, sections were fixed by incubating with fresh 4% PFA for 30 minutes diluted in PBS. After that, samples were washed 2x with fresh PBS and dehydrated with EtOH. The EtOH solutions were prepared immediately before the dehydration and were diluted in Ambion™ DEPC-treated water (Invitrogen, #AM9920) When the EtOH was completely evaporated, a hydrophobic barrier was created with the ImmedgeTM hydrophobic barrier pen (Vector Laboratories, #H-4000). Then, samples were treated with RNAscope® Protease IV for 30 minutes at RT. Next, sections were washed 2x with PBS and were hybridized in the HybEZTM Oven for 2 hours at 40°C with the RNA probes Mouse TNF-α Atto-550, HDV genome Alexa-Fluor 488, and HDV antigenome Atto-647. After that, the slides were washed 2x with 1X Wash Buffer and were incubated with the RNAscope® Detection Reagents to amplify the hybridization signals. Finally, the liver sections were incubated for 30 seconds with DAPI and mounted with ProLong™ Gold Antifade Mountant solution (Thermo Fisher, #P10144). For ISH combined with F4/80 IF HDV antigenome probe was removed, and after the last incubation with the RNAscope® Detection Reagents, anti-F4/80 antibody (Table 1) was added diluted 1:10,000 and incubated overnight at 4°C. Then slides were washed 3 times with PBS and incubated with anti-rat antibody for 30 min diluted 1:200, followed by an anti-rabbit Alexa 647 for 1h at 1:200. Finally, the liver sections were incubated for 30 seconds with DAPI and mounted with ProLong™ Gold Antifade Mountant solution (ThermoFisher, #P10144). Fluorescent Images were acquired with the LSM 880 (Zeiss, Jena, Germany) laser-scanning confocal microscope and the Vectra® Polaris™ Automated Quantitative Pathology Imaging System (Perkin Elmer). Then, the obtained images were quantified with an ImageJ Plugin developed by the Imaging Platform at CIMA.

### CRISPR/Cas9 vectors and gRNA design

The pX602-AAV-TBG::NLS-SaCas9-NLS-HA-OLLAS-bGHpA;U6::BsaI-sgRNA plasmid that contains the Staphylococcus aureus Cas9 (SaCas9) expressed under the TBG promoter, the sgRNA under U6 promoter and ITR sequences for AAV vector production was a gift from Feng Zhang (Addgene plasmid # 61593). sgRNAs targeting exonic regions of the Ripk1 gene were designed and selected using Benchling software (www.benchling.com). The selected 21-nt sequences upstream of the 5’-NNGRRT-3’ PAM sequence of SaCas9 are shown in Table 2. Annealed oligonucleotides coding for the guide RNA sequences (Sigma) were cloned into BsaI site of pX602 vector using standard molecular cloning techniques.

### Recombinant AAV production

The AAV genomes were packaged in AAV serotype 8-capsids (AAV8) as previously described [16]. Briefly, for each production, the AAV shuttle vector and the packaging plasmid pDP8.ape (Plasmid factory) were co-transfected into HEK293T cells. The cells and supernatants were harvested 72 h upon transfection and virus was released from the cells by three rounds of freeze–thawing. Crude lysate from all batches was then treated with DNAse and RNAse (0.1 mg per p150 culture dish) for 1 h at 37 °C and then kept at −80 °C until purification. Purification of crude lysate was performed by ultracentrifugation in Optiprep Density Gradient Medium-Iodixanol (SigmaAldrich). Thereafter, iodioxanol was removed and the batches concentrated by passage through Amicon Ultra-15 tubes (Ultracel-100K; Merck Millipore). AAV8 vector without sgRNA was also produced as nonediting control. For virus titration, viral DNA was isolated using the High Pure Viral Nucleic Acid kit (Roche Applied Science). Viral titers in terms of viral genome per milliliter (vg/mL) were determined by qPCR (Applied Biosystems) using SaCas9 or AAT specific primers (Table 2).

### Serum ALT levels

Alanine aminotransferase (ALT) serum levels were analyzed with a Hitachi Automatic Analyzer (Boehringer Mannheim, Indianapolis, IN).

### Histology and immunohistochemistry (IHC)

Liver sections were fixed with 4% paraformaldehyde (PFA), embedded in paraffin, sectioned (3 μm), and stained with hematoxylin and eosin and were mounted and analyzed by light microscopy for histological evaluation. For IHC all reactions required antigen retrieval, 30 min at 95 °C in 0,01 M Tris-1 mM EDTA pH 9. Incubations with primary antibodies at their optimal dilutions were performed overnight at 4°C. Primary antibodies used for the IHC were: F4/80 (BioLegend 123102, 1:400), cleaved Caspase-3 (activated Caspase 3 (a-Casp3)) (Cell Signaling 9661, 1:200) (Table 1). For HDAg analysis, patient serum with anti-HDAg reactivity was employed as primary antibody (1:10,000). After rinsing in TBS-T, the sections were incubated with the corresponding secondary antibodies for 30 min at RT. Peroxidase activity was revealed using DAB+ and sections were lightly counterstained with Harris hematoxylin. Finally, slides were dehydrated in graded series of ethanol, cleared in xylene and mounted with Eukitt (Labolan, 28500). Image acquisition was performed on an Aperio CS2 slide scanner using ScanScope Software (Leica Biosystems). All images were stored in uncompressed 24-bit color TIFF format and image analysis was performed using a plugin developed for Fiji, ImageJ (NIH, Bethesda, MD).

Samples from patients included in the study were provided by the Biobank of the University of Navarra and were processed following standard operating procedures approved by the Ethical and Scientific Committees (Reference: 2020-011).

### Protein extraction and Western Blotting

Liver samples were lysed in RIPA buffer (20 mM Tris-HCL pH 7.5, 150 mM NaCl, 1% NaDeoxycholate, 1% Triton X-100 and 1% SDS) supplemented with protease inhibitors. Protein concentration was determined by Pierce™ BCA Protein Assay Kit (Thermo Scientific, US) according to the manufacturer’s recommendations. Western blot was performed using mouse anti-GAPDH (G8795, Sigma-Aldrich, 1:5000), and rabbit anti-RIPK1 (#3493, Cell Signalling, 1:1000) in PBS-T 5% powdered milk (Table 1). This was followed by horse anti-mouse IgG (7076P2, Cell Signalling, 1:5000) or goat anti-rabbit IgG (7074S, Cell Signalling, 1:5,000), respectively. SuperSignal® West Pico Chemiluminescent Substrate (Thermo Scientific) was used to detect expression.

### Statistical analysis

Statistical analysis was performed using GraphPad Prism 7.0 software. The data are presented as mean values ± standard deviation. Differences in ALT levels and % of stained tissue area between groups were analyzed with one-way or two-way ANOVA followed by Bonferroni multiple-comparison test or Kruskal Wallis test followed by Dunns’ multiple comparison test. Pearson correlation analysis was performed, and the correlation coefficient calculated. (Significance *P< 0.05, **P < 0.01, ***P < 0.001, ****P<0.0001). Mann

## Acknowledgments

We particularly acknowledge the patients for their participation and the Biobank of the University of Navarra for its collaboration. We thank Professor John Taylor, Professor Frank Chisari and Professor Feng Zhang for providing us with essential reagents for our study. We are grateful to Elena Ciordia, Alberto Espinal, and CIFA staff for animal care and vivarium management and to Laura Guembe and the imaging department at CIMA for technical assistance.

## Author contributions to manuscript

Gracian Camps, Sheila Maestro, Carla Usai, acquisition, analysis, statistical analysis and interpretation of data. Laura Torella design of CRISPR-Cas9 gene editing systems. Ana Aldaz, quantitative and qualitative analysis of the genetic modifications after gene editing. Lester Suarez design and construction of the recombinant AAV-HDV vector. Cristina Olague and Africa Vales animal manipulation and technical support. Anne Montfort, Bruno Ségui, provide key experimental tools. Gracián Camps drafting of the manuscript. Rafael Aldabe and Gloria Gonzalez-Aseguinolaza study concept, design, and analysis, funding acquisition, and preparation of the final version of the manuscript.

## Abbreviations

AAV: adenoassociated virus
a-Casp3: activated caspase 3
ALT: alanine aminotransferase
Casp3: Caspase 3
CRISPR: Clustered Regularly Interspaced Short Palindromic Repeats
CRISPR-Cas9: CRISPR and Cas9 nuclease
HBV: hepatitis B virus
HBsAg: hepatitis B surface antigen
HCC: hepatocellular carcinoma
HDV: hepatitis D virus
HDAg: HDV antigen
IF: Immunofluorescence
IFN: interferon
IHC: immunohistochemistry
ISH: In situ hybridization
LPS: lipopolysaccharides
MAVS: mitochondrial antiviral signaling protein
MHV3: murine hepatitis virus type 3
MLKL: Mixed lineage kinase domain like protein
Nec-1: necrostatin 1
NK: natural killer
rAAV: recombinant AAV
Rag1: recombination activating gene 1
RIPK: receptor-interacting serine/threonine-protein kinase
TNF: tumor necrosis factor

## Supplementary figures

**S1 Fig.**
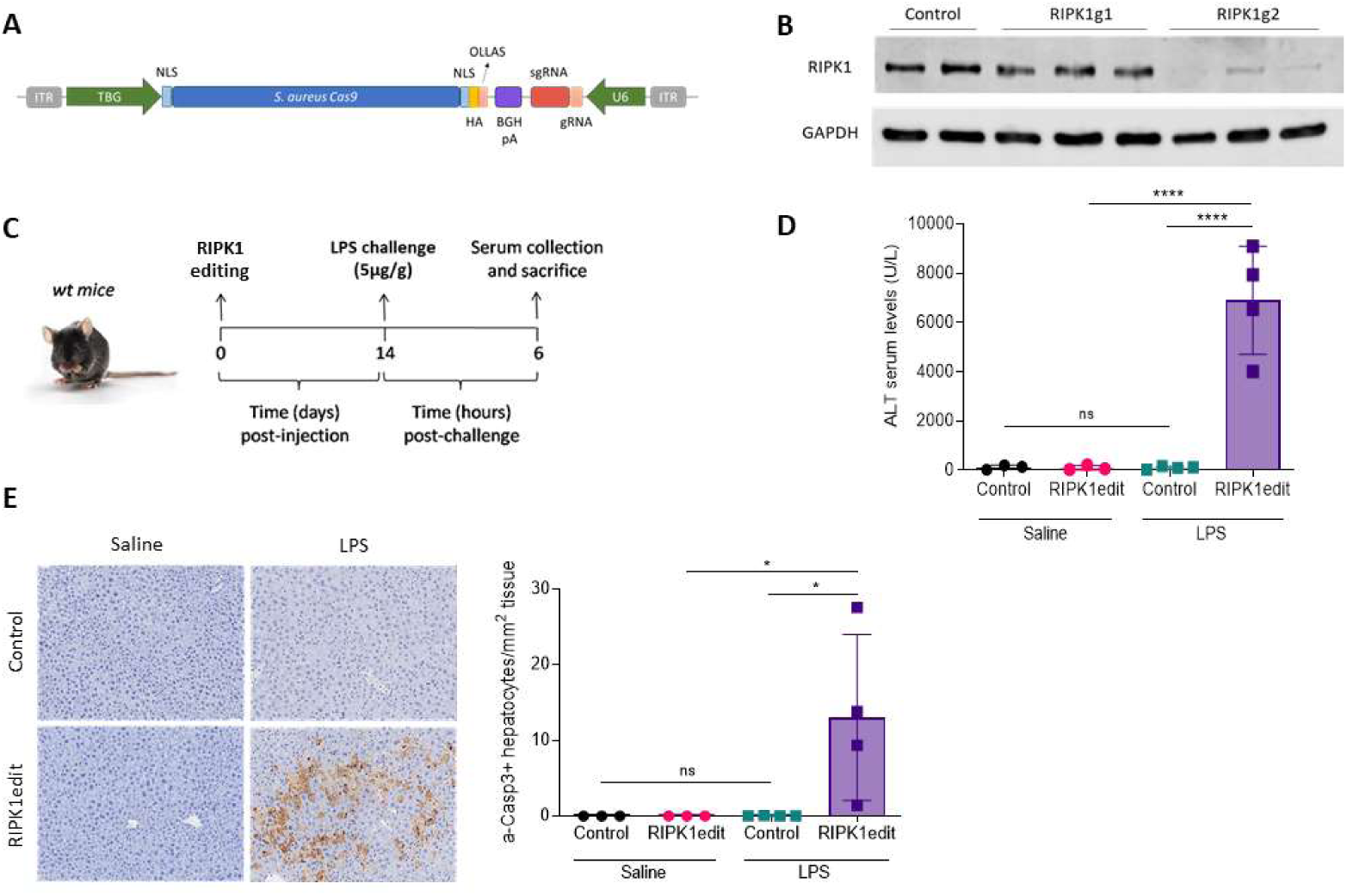
(A) Schematic representation of the recombinant AAV genome carrying the Staphylococcus aureus Cas9 protein flanked by two nuclear localization signals (NLS) and fused to the OLLAS tag under the control of a liver specific promoter TBG (Thioglobulin promoter) and the guide RNA (compose by the sgRNA and the gRNA) under the control of U1 promoter. (B) C57BL/6 mice received 10^11^ gc of AAV-SaCas9-RIPK1g1 or AAV-SaCas9-RIPK1g2 and 30 hours later animals were sacrificed and RIPK1 expression was analysed by western blot in liver extracts. (C). Schematic representation of the experimental procedure, 6-8-week-old C57BL/6 wt mice were intravenously injected with 10^11^ genome copies (gc) of AAV-SaCas9-RIPK1g2 (RIPK1edit) or an AAV expressing SaCas9 without guide (control) and 14 days later animals were challenged with LPS at a dose of 5 µg/gr and sacrificed 6 hours later. Liver damage was analyzed by (D) quantification of serum ALT levels (U/L) and (D) quantification of a-Casp3+ hepatocytes/area after Immunohistochemistry (IHC) analysis. Statistical analysis was performed by one-way ANOVA followed by Bonferroni multiple-comparison test. p <0.05 (*), p <0.0001 (****), ns = non-significant.

**S2 Fig.**
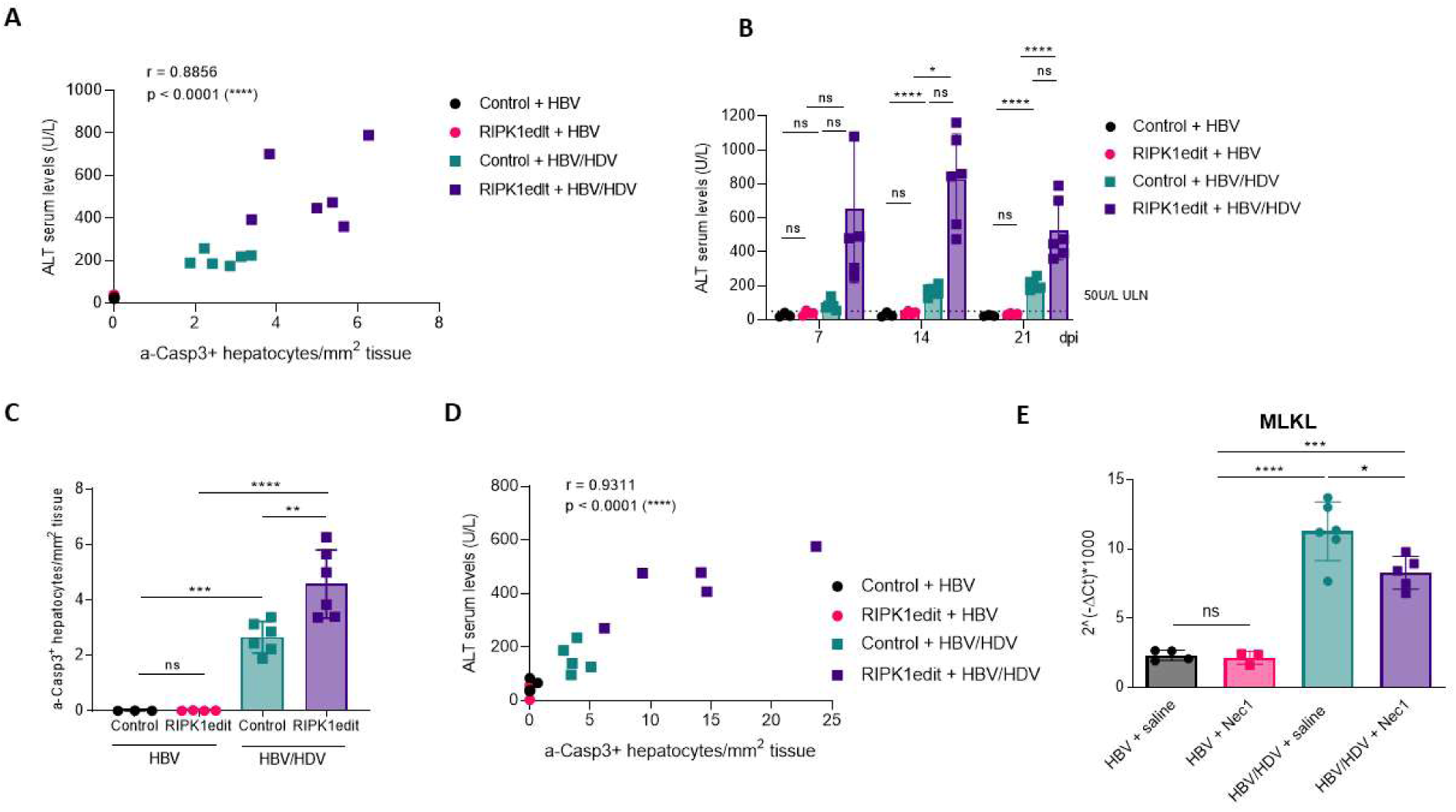
(A) Correlation between ALT levels and a-Casp3 positive hepatocytes in wt mice treated as described in fig 2A. (B-D) C57BL/6 Rag1 KO mice were treated as described in Fig 2A and liver damage was analyzed by (B) quantification of serum ALT levels (U/L) and (C) quantification of a-Casp3+ hepatocytes/area after Immunohistochemistry (IHC) analysis. (D) Correlation between ALT levels and a-Casp3 positive hepatocytes. (E) MLKL expression levels was analyzed in the liver of mice treated daily with a dose of 2.5 mg/kg Nec1 of saline and that were previously injected with injected AAV-HBV (HBV) or AAV-HBV/HDV (HBV/HDV). Statistical analysis was performed by one-way ANOVA followed by Bonferroni multiple-comparison test. p <0.05 (*), p <0.01 (**), p <0.001 (***), p <0.0001 (****) and ns = non-significant.

**S3 Fig.**
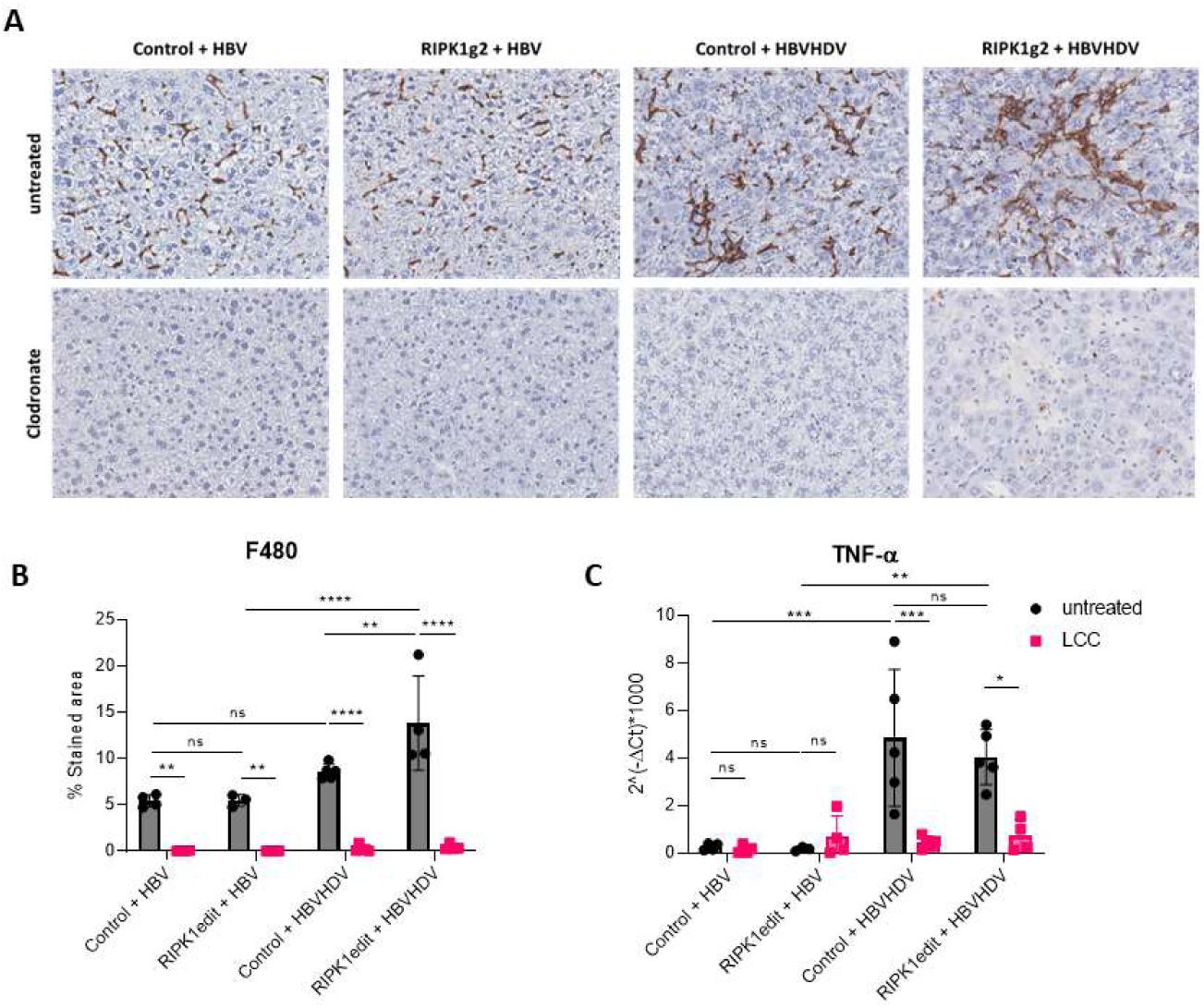
Analysis of macrophage depletion by anti-F4/80 immunohistochemistry in mice receiving saline or clodronate loaded liposomes previously treated as described in Fig 2A. (A) representative images from the different groups of animals (B) quantitative analysis. (C) At sacrifice TNF-α expression was analyzed in the liver of mice by RNA retrotranscription followed by quantitative PCR.

